# Predicting proprioceptive cortical anatomy and neural coding with topographic autoencoders

**DOI:** 10.1101/2021.12.10.472161

**Authors:** Max Grogan, Kyle P. Blum, Yufei Wu, J. Alex Harston, Lee E. Miller, A. Aldo Faisal

**Affiliations:** Department of Bioengineering, Imperial College London, London, UK; Department of Physiology, Northwestern University, Illinois, USA; Institute of Artificial & Human Intelligence, University of Bayreuth, Bayreuth, Germany

**Author notes:** These authors contributed equally to the work (Max Grogan, Kyle P. Blum, Yufei Wu).

## Abstract

Proprioception is one of the least understood senses, yet fundamental for the control of movement. Even basic questions of how limb pose is represented in the somatosensory cortex are unclear. We developed a topographic variational autoencoder with lateral connectivity (topo-VAE) to compute a putative cortical map from a large set of natural movement data. Although not fitted to neural data, our model reproduces two sets of observations from monkey centre-out reaching: 1. The shape and velocity dependence of proprioceptive receptive fields in hand-centered coordinates despite the model having no knowledge of arm kinematics or hand coordinate systems. 2. The distribution of neuronal preferred directions (PDs) recorded from multi-electrode arrays. The model makes several testable predictions: 1. Encoding across the cortex has a blob-and-pinwheel-type geometry of PDs. 2. Few neurons will encode just a single joint. Our model provides a principled basis for understanding of sensorimotor representations, and the theoretical basis of neural manifolds, with applications to the restoration of sensory feedback in brain-computer interfaces and the control of humanoid robots.

**Author Summary:** It is well established that proprioception is essential for effective for motor control, yet our understanding of proprioceptive coding in somatosensory cortex is far behind that of more established sensory modalities such as vision and touch. Here, we use unsupervised learning of a deep neural network imposed with biological constraints to reproduce coding properties of proprioceptive neurons in area 2 of primary somatosensory cortex. With this model, we demonstrate that the tendency for area 2 neurons in close physical proximity to share similar directional tuning can be explained by local topographic organisation driven by Mexican hat lateral effects, a phenomenon that is otherwise unobservable with the spatial resolution of available electrode microarray recordings. This provides, to the best of our knowledge, the first evidence of local topographic organisation in area 2 proprioceptive neurons. We also predict that the structure of proprioceptive receptive fields are typically multi-joint rather than single-joint in nature and provide, to the best of our knowledge, and find that our model best reproduces the tuning properties of neural data when training on natural kinematic recordings, rather than those from a stereotyped task, underlining the importance of training on data that reflect the natural distribution of sensory stimuli.

## Introduction

Somatosensation includes the familiar sense of touch, provided by receptors in the skin, and proprioception, the much less consciously perceived sense that informs us about the pose, motion and associated forces acting on our limbs. While the former has received much scientific attention, proprioception is often overlooked, yet this modality of sensory feedback is essential for our ability to plan, control and adapt movements. In engineering, the control of robotic movement would be impossible if the controller did not know the location of its actuators; correspondingly in human motor control, optimal feedback control theory is the preeminent explanation for the computations underlying limb control [1,2]. Moreover, individuals with proprioceptive neurological deficits, such as patient IW, have profound motor deficits even in the presence of vision and an intact motor system [3,4]. Similarly, recent major developments in neuroprosthetics are centred on restoring sensory feedback as well as limb motion through bi-directional interfaces [5] and will likely require not only touch but also proprioceptive feedback to restore functional capability [6]. Thus, understanding proprioceptive encoding is essential for restoration of both motor action and sensory function in clinical rehabilitation [7].

Unlike hearing or vision, proprioceptive afferent pathways originate not from a single organ but from diverse families of mechanoreceptors within muscles, tendons, joints, and the skin itself [8]. Proprioceptive information ascends within the dorsal column pathway through the dorsal root ganglia, the dorsal column nuclei, and thalamus before arriving in the primary somatosensory cortex (S1). Crucially, neurons are somatotopically organised throughout this pathway, meaning that neighbouring neurons encode stimuli from closely related parts of the body. This gives rise to the sensory homunculus, which results from the ordered mapping of tactile representations of the body’s surface across the cortical surface [9].

Although proprioception is generally acknowledged to be critical to motor behaviour, the corresponding proprioceptive maps – particularly that of primate area 3a, but also the mixed modality area 2 – are much less distinct and well understood than those of the touch (areas 1 and 3b). We know that the proprioceptive and tactile systems often encode overlapping information, e.g. mechanoreceptors in our skin and interosseous membranes respond to deformation and vibration, contributing to the sense of body position and movement [3]. Indeed, the firing rates of somatosensory neurons with cutaneous receptive fields in area 2 can be used to decode limb movement as accurately as those with muscle fields [10]. While proprioceptive cortical areas are critical for our ability to generate goal-directed complex behaviour it is unclear what properties of proprioceptive cortical coding facilitate this capability. Crucially, we lack an accepted hypothesis about the computational principles that drive the mapping of proprioceptive arm representations onto the cortex.

The limited nature of our understanding of proprioceptive neural representations in the brain has two major causes. First is the difficulty recording with many electrodes from sulcal proprioceptive areas in primates [11], and second is the difficulty delivering independent proprioceptive stimuli in comparison to other senses, such as vision [12] and touch [13]. To overcome these limitations, we combined computational modelling and natural movement kinematics data to test hypotheses of organisational mechanisms that are currently beyond the capability of experimental recording techniques. This combination of experiment and theory has been useful for explaining population-level coding in other sensory modalities, such as vision [14], olfaction, and touch [15,16], but has only recently begun to be applied to proprioception [17]. Much of the computational theoretical work on other senses has either focused on predicting specific neural coding features from natural sensory statistics [15,18], or has ignored the details of neural coding and focused on the spatial distribution of stimulus representation across the cortex [19].

Our chosen system was the proprioception of the right arm (Fig. 1.A) for which we had collected natural behavioural data from daily life in humans (food preparation, eating, etc; Fig. 1.B and Supplemental Fig. S1) as well as constrained, planar centre-out reaching in both human and monkeys (Fig. 1.C). This data describes the joint angle kinematics of the of the body over time and thus stands in for the proprioceptive sensory state. We aimed to relate kinematics to proprioceptive encoding with modelling and comparisons to existing single-unit recordings from monkeys. We formulated a novel computational model that predicts both neural coding of single neurons and spatial organisation of these neurons across the cortex (Fig. 1.D-G). We chose a small set of general computational elements that reflect principles and mechanisms found in sensory systems but have not been previously unified, which are as follows: Models should 1) use the information maximisation principle, which postulates that efficient sensory representations in the brain reflect natural sensory statistics, and implies that they are essential in shaping neural representations, 2) be stochastic, generative, and decodable, to reflect the natural variability in the data, produce neural activity to mimic that of the biological system, and offer the means to reconstruct the relevant original sensory input from its output, 3) implement (in Marr’s sense [20]) neural computations that are performed by locally interacting neurons through synaptic interactions, rather than by an abstract computational machine.

**Figure 1:**
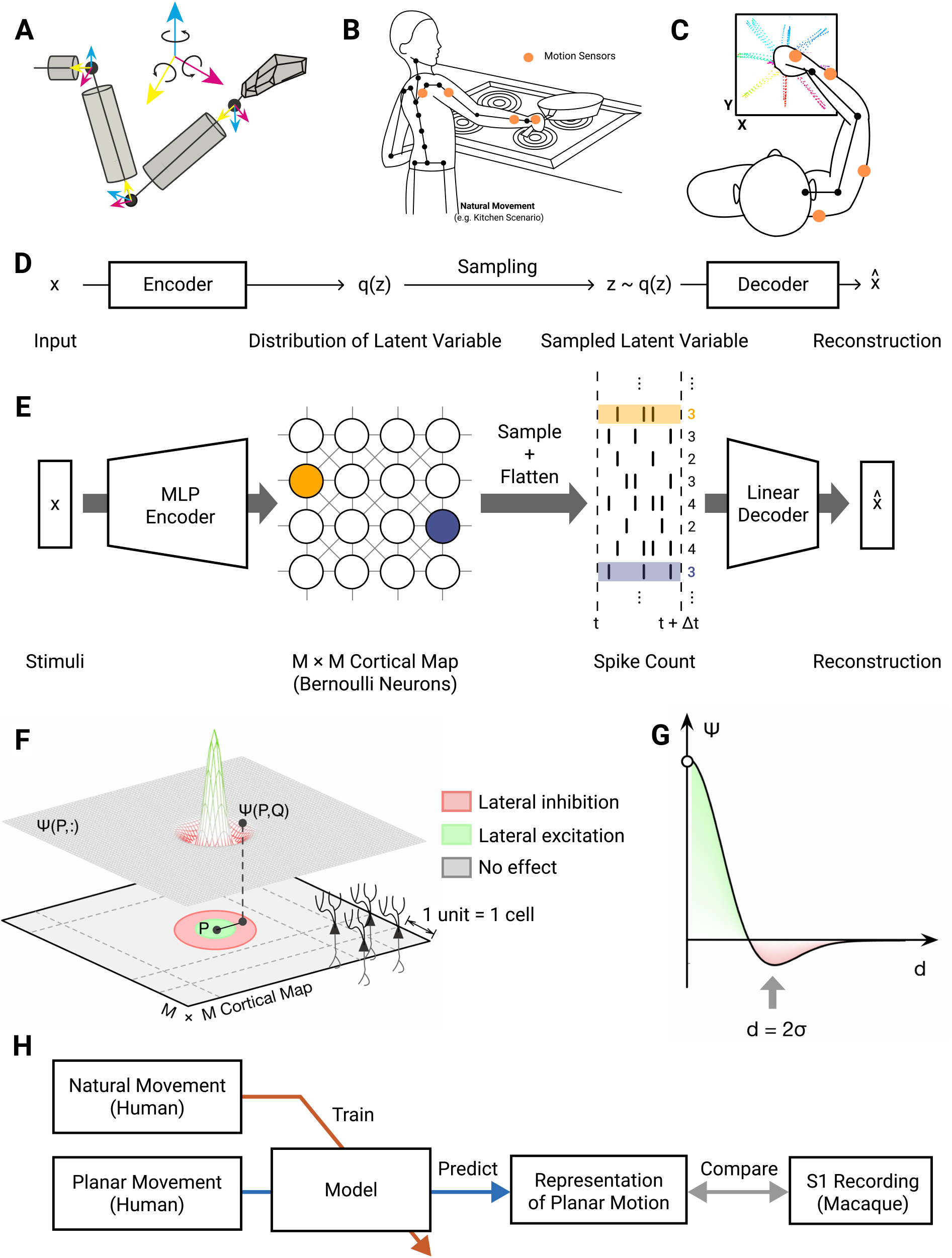
Task Context and model architecture. (**A**) 9-dimensional arm movement data acquired from motion capture suit is represented by Euler angles between body segments, following the ISB Euler angle extractions [25] in a ZXY coordinate. **(B)** Illustrative example of movements carried out in the natural scenario (cf. Fig. S1 for further examples). **(C)** Illustrative example of planar centre-out reaching movements. **(D)** General architecture of a VAE: Inputs are encoded as a latent distribution, from which samples are decoded to reconstruct the original input. **(E)** Topo-VAE architecture. Our cortical neural representation is modelled by an 80×80 cortical grid of artificial neurons in the latent layer q(z). In contrast to conventional VAEs which model these latent neurons as multi-dimensional Gaussian random variables, we used Poisson random variables. Input stimuli drive these model neurons via a multi-layer perception (MLP) encoder network (2 layers of 500 neurons each). A linear decoder is applied to the spike counts emitted from the cortical map within a time interval Δt, to reconstruct the sensory input stimuli. **(F)** To embed the model neurons in a cortex-like topographic context we use a Mexican-hat lateral effect between the latent layer neurons. **(G)** This interaction Ψ(P,Q) is a function of the Euclidean distance d = ∥ P-Q ∥ between a pair of neurons P,Q and is characterised by a length scale σ. Nearby neurons are excited, intermediate-range neurons are inhibited and there is no effect on distant neurons. **(H)** Flow of natural and planar movement data (joint angle velocities) in this work.

In the context of proprioception, these principles require that our model should learn how to translate proprioceptive inputs from movements of the body into a latent representation of proprioception (which is read out as neural firing). In addition to performing efficient feature learning of a spike-based code, the model should have minimal human induction bias, except for specific characteristics relating to the principles laid out above. By training the model on natural movement stimuli, it should learn representations that can maximise information about the natural environment with the limited bandwidth of neurons that is available at any given encoding step, thereby implementing efficient coding [21–23]. Given these requirements, we chose to use an unsupervised deep neural network model, a variational autoencoder (VAE) [24] (Fig. 1.D) to perform efficient feature learning of proprioceptive stimuli. The primary training objective of an autoencoder is to reconstruct the input stimuli, enabling the model to learn in an unsupervised manner. Furthermore, the bottleneck of the VAE model parameterises a latent Bernoulli distribution to imitate the activity of spiking neurons.

In following with our third outlined principle, and the observation that spatial organization is relevant to cortical coding, we introduce a 2-dimensional structure to the latent (“bottleneck”) layer of the VAE, representing a simplified proprioceptive cortex (Fig. 1.E). A conventional “vanilla” VAE model will capture the properties of encoded sensory features but would be devoid of any of the spatial properties of cortical neurons that are critical for understanding how sensory stimuli are represented across the cortical surface. Incorporating spatial relationships in neural coding models has not been well investigated, yet anatomical structure and function in neural computation are fundamentally linked by biophysical constraints [26]. We therefore also implemented a topographical mechanism, using a lateral interaction term between the units in the VAE’s latent layer (see Methods for detailed motivation and mathematical description; Fig. 1.F). This term corresponds biologically to a spatial distribution of short-range excitatory and longer-range inhibitory synaptic interactions between neurons (Fig. 1.G).

We call our model the “topo-VAE” and compared it with existing recordings from area 2 of somatosensory cortex. It could equally well be applied to other proprioceptive areas such as 3a or even 5. Models with its basic form could be applied to other sensory modalities, including touch, vision, and hearing. Here, the approach has enabled us to uncover novel insights into proprioceptive coding by linking anatomical structure and function so that we can test our model’s predictions using the sparse set of obtainable data.

## Results

We developed a novel model of cortical sensory representation (see Methods for details) that predicts both function and structure – the topo-VAE model. We trained, tested and validated our model as follows (Fig. 1.H): We trained our topo-VAE model on natural daily human arm movement kinematics and explored the emergent proprioceptive representations in its cortical layer (i.e. the latent layer of the topo-VAE). After training, we used the model to generate neural responses by providing it kinematic data from a centre-out reaching task performed by human subjects. We then compared the properties of these simulated neural activities (i.e. the generated spike trains) to those of S1 neurons which were previously recorded from monkeys performing a planar centre-out reaching task. The human and monkey reaching data were rescaled so that the biomechanical differences between human and monkey arm movements are kept small. We characterised the proprioceptive neural representations at two levels of description: first, we considered the spatial tuning curves of individual neurons and second, and second, we considered the organisation of tuning preference maps across our model’s cortical surface.

We used three sets of measurements to summarise the neural coding properties of S1 neurons for this centre-out task: 1) the preferred direction (PD) of single-neuron firing rates, 2) features of the tuning surface defined by firing rate modulation as a function of both direction and speed of movement, and 3) the overall distribution of firing rates throughout the behaviour. First, we analysed PDs of neurons in the recordings and in our modelling (Fig. 2). The PD refers to the movement direction of the hand (when performing the centre-out reaching task) in which a neuron was most active. We found that generally, modelled neurons exhibited PD tuning surfaces that were similar in appearance to those of recorded neurons (Fig. 2.A,B respectively). At the population level, while there was a statistically significant difference (p<0.005; Kolmogorov-Smirnov test) between the distribution of PDs of modelled and recorded neurons (Fig. 2.C), the distributions were similar in that they were both bimodal (Supplemental Fig. 2), with a bias towards the 120°/300° reach axis (Fig. 2.D). Previous experimental studies have suggested such coding biases are a natural consequence of musculoskeletal constraints in the arm [27,28], which cause the natural distribution of right arm reach movements to favour the 120°/300° reach axis. Under the efficient coding hypothesis, this becomes reflected in the coding properties of the sensory neurons. To test this theory, we trained a second model on stereotyped movement stimuli from the centre-out reach task only. We found that this new model was able to reconstruct natural movement data almost as well as the natural-trained model (0.80 ±0.01 vs 0.96 ±0.008, planar-trained vs natural-trained; R^2^ scores; p<0.005; Wilcoxon) and produced a PD distribution that was also comparable to the recorded neuron distribution (0.241 vs 0.248, planar-trained vs natural-trained; Jensen-Shannon divergence). However, unlike the natural-trained model, we observed a reach axis bias that was perpendicular to that of recorded neurons (Fig. 2.D,E). This suggests that certain features of proprioceptive coding are only emergent when the coding is conditioned on the natural distribution of stimuli.

**Figure 2.**
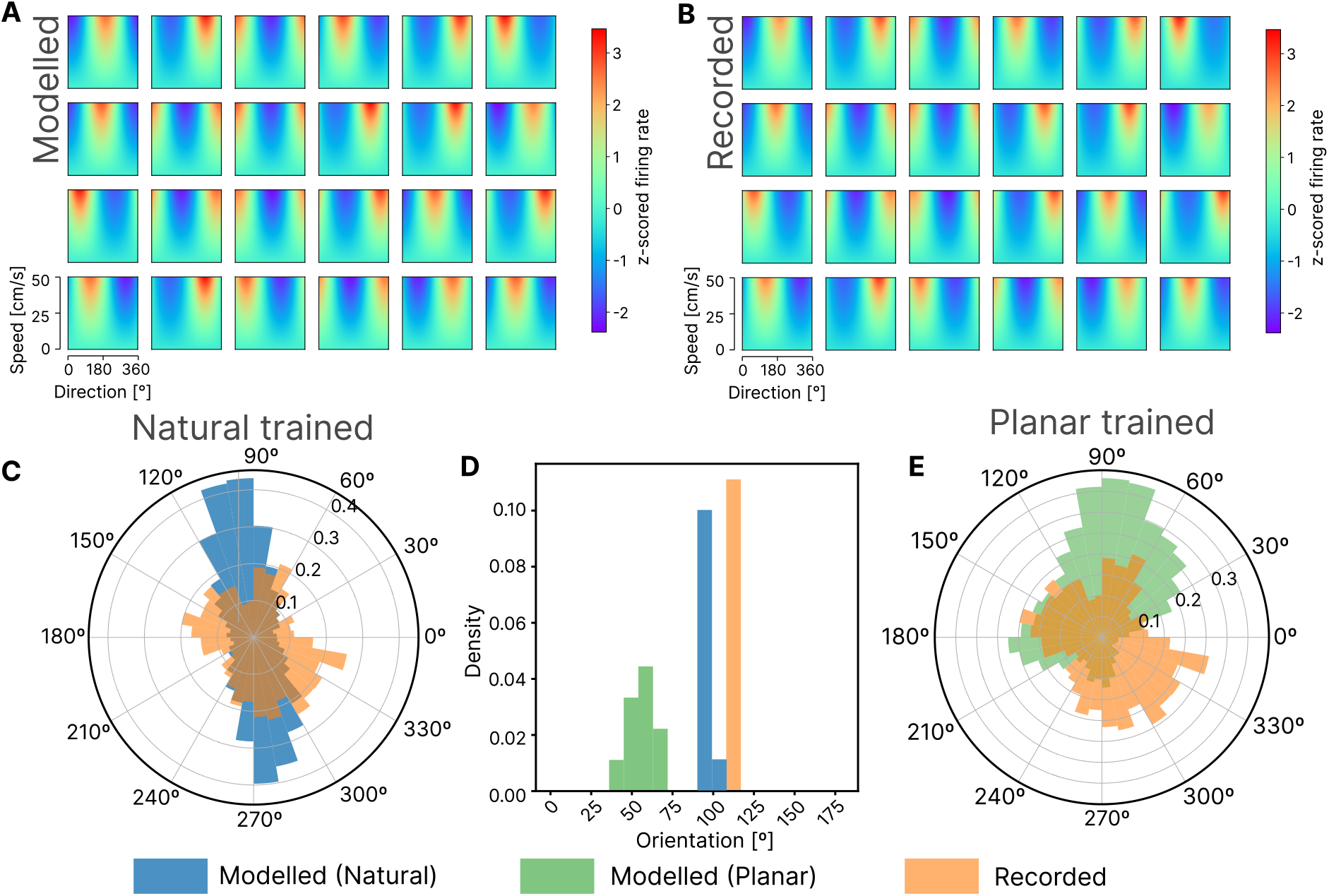
Comparing tuning of modelled and recorded neurons. (**A)** Tuning maps of example area 2 neurons during planar movements. **(B)** Tuning maps from example neurons observed in the latent layer of the topo-VAE under planar movements, matched to example neurons in A. (**C)** Circular density histograms of preferred directions in our topo-VAE model trained on natural data (n=6400) (blue) and neurons recorded from area 2 of 3 monkeys (n=383) (orange). Note, that the neural models were not fitted to monkey neural data, but directly predicted from the statistics of natural human body kinematics. **(D)** Frequency histogram of mode bias in PD distributions for topo-VAE models trained on natural data (blue) and stereotyped centre-out reach data (n=10 for each condition) vs mode bias of PD distribution in recorded neurons (Orange line). **(E)** Circular density histograms of preferred directions in our topo-VAE trained on stereotyped centre-out reach data (n=6400) (green) and neurons recorded from area 2 of 3 monkeys (n=383) (orange).

While preferred direction is an informative neural correlate, we looked to better compare tuning properties by extracting geometric features from the tuning surfaces of modelled and recorded neurons and evaluating how different model properties affect similarity. Two such features were compared (Fig. 3.A): Half-peak widths measure how sharp the tuning of a neuron is, where smaller half-peak widths indicate sharper tuning, and velocity gradients measure how sharply the firing rate of a neuron changes as a function of the endpoint velocity in the direction of tuning (note that the velocity gradient measurements were adjusted to control for differences in mean firing rate between modelled and recorded neurons). While there was a statistically significant difference between the feature distributions of modelled and recorded neurons (p<0.005 for both features and all model variations; Kolmogorov-Smirnov test), Jensen-Shannon divergence between distributions was minimised (Fig. 3.B,C) when the model included lateral effects and was trained on natural movement data. This would suggest that both model constraints are important for reproducing the tuning properties of recorded neurons. One concern was that the distributions of tuning curve geometry between modelled and recorded neuron populations were similar only because of the Poisson GLM fitting process –-perhaps fitting a Poisson GLM to any random data would produce distributions like these. To verify that this was not the case, we shuffled the firing rate data from modelled neurons to remove any correlation between endpoint velocities and firing rates and trained an additional Poisson GLM on this data. The shuffling produced a very different pair of distributions for the geometric features, with flat, poorly tuned curves – suggesting that any similarities we saw between the original unshuffled curves were in fact due to similarities in the underlying structure of the modelled and recorded neuron data. This provides an empirical lower bound of the similarity between feature distributions, and would suggest that similarity between the distributions of modelled and recorded neurons is not trivially the result of using a GLM (Fig. 3.B,C).

**Figure 3.**
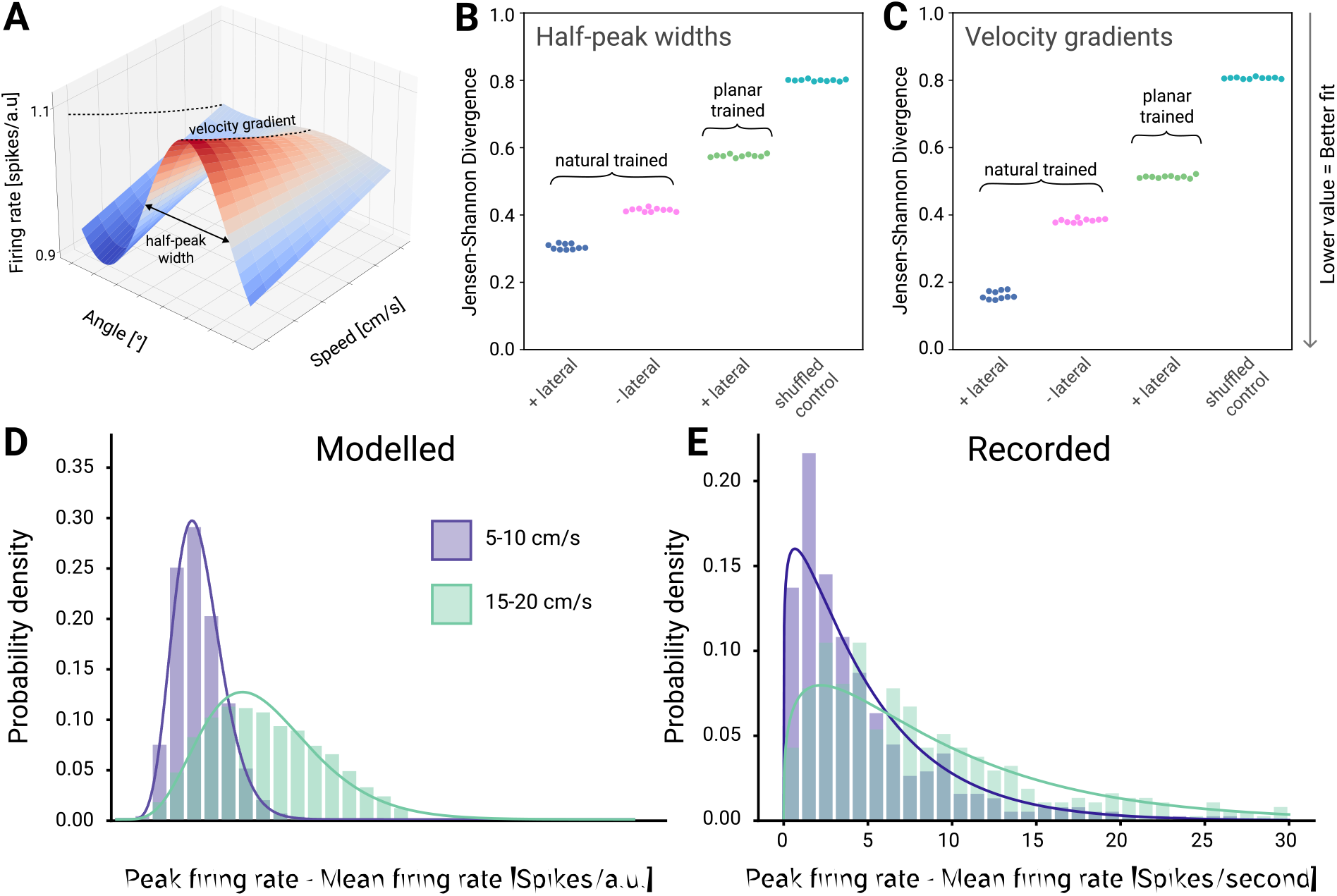
Comparing tuning surface features and firing rate distributions between modelled and recorded neurons. (**A)** Tuning surface from a single modelled neuron, illustrating two measures we computed to summarise neuronal modulation with velocity: half-peak width of spatial tuning (at maximum end-point speed) and velocity gradient. (**B)** Jensen-Shannon divergence between the distributions of half-peak widths in modelled neurons trained under different conditions (with (+) and without (-) lateral effects, trained on natural data, trained on planar data) (6400 neurons, 10 models per condition) and recorded neurons (383 neurons). The shuffled control is the same as the natural trained model with lateral effects, except the neural activity is shuffled prior to training the Poisson-GLM in the tuning surface calculation step. **(C)** Equivalent plot to (B) for velocity gradient distributions. (**D)** Firing rate modulation (peak rate minus mean rate) by endpoint-velocity across all reach directions in modelled (6400 neurons) and (**E)** in recorded neurons (383 neurons). Histograms represent actual distributions, whereas lines represent fitted gamma distributions (R^2^>0.95 for both; Kolmogorov-Smirnov test).

Finally, we examined endpoint-velocity modulation of firing rate distributions, which were well fit by Gamma curves for both modelled (Fig. 3.D) and recorded neurons (Fig. 3.E; R^2^ > 0.95 for equal numbers of neurons). While the topo-VAE generates spike count distributions over an arbitrary time window (cf. Fig. 3.D, x-axis), the functional form and relative change of the distributions with altered velocity are not arbitrary. Therefore, we compared the distributions at different endpoint velocities and found that they differed significantly in both modelled and recorded neurons (p<0.005; two-sided Wilcoxon signed-rank test), indicating velocity dependence in both the modelled and recorded neurons.

In summary, modelled neurons in the topo-VAE exhibit various tuning properties such as preferred direction and endpoint-velocity modulation of firing rates which are structurally similar to those of recorded neurons. While the population statistics of these tuning properties differ significantly between modelled recorded neurons, the combined lateral effects that constrain our model and the training data that reflect the natural distribution of arm movement, increase the similarity, all without fitting our model to any neural data.

Having demonstrated similarity in the tuning statistics between modelled and recorded neurons, we next compared the spatial relationship of PDs in our model neurons to those in the monkey brain within the constraints of the 400 μm spacing of the recording electrodes (Fig. 4). While neurons recorded at two adjacent electrodes corresponded to a distance that was substantially beyond the effective range of the Mexican hat distance function, many electrodes recorded data from more than one distinguishable neuron. The distance between these neurons was within the local neighbourhood of neurons in our model. We could thus compare the similarity of neural coding properties (specifically, the PDs) between neurons recorded on the same electrode to the similarity between neurons on different electrodes (cf Fig. 4.A). In modelled neurons, same-electrode PD differences followed an exponential distribution, such that the probability that two neurons had similar (within 30 degrees) PDs was much higher than chance (p=0.77; two-sided Wilcoxon signed-rank test; Fig. 4.C., blue histogram). Conversely, the probability that two neurons on the same electrode had nearly opposite PDs (between 150° and 180°) was very low (p=0.03; two-sided Wilcoxon signed-rank test; Fig. 4.C orange histogram). We then ran the same analysis on recorded neurons and found that the PD difference distributions appeared similar to those predicted by the model (Fig. 4.C). Given this similarity, we use the topo-VAE model to directly predict the spatial organization of PD tuning across the proprioceptive cortex, which the 400um electrode spacing lacks the adequate spatial resolution to observe. The model predicts that PDs are clustered in blob-like structures of similar PDs with boundary regions where the PDs rotate smoothly toward neighbouring directions (Fig. 4.D). This arrangement is interrupted by small, sparsely spaced “pinwheel” regions, representing all PDs in a small neighbourhood, which is consistent with same-electrode similarity of Fig. 4.B,C.

**Figure 4.**
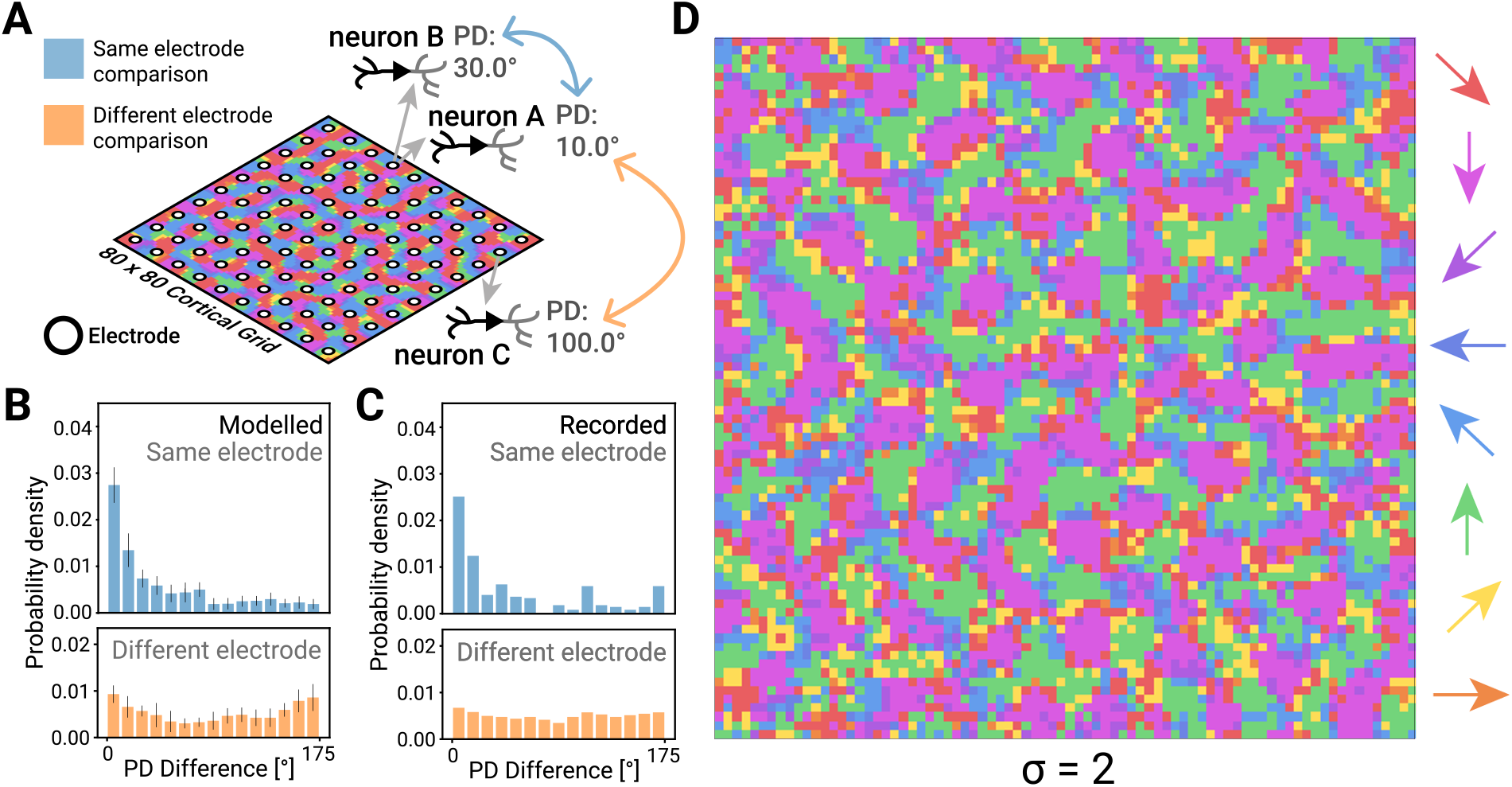
Predicting the topography of proprioceptive cortex. (**A)** Illustration of pairwise comparisons of preferred directions recorded on same (blue) and different (orange) electrodes, performed in recorded and modelled neurons. Neuron A and B are recorded from the same electrode, Neuron C is recorded from a distant electrode. **(B)** Distribution of PD difference for same electrode (blue histograms) and different electrode (orange histograms) comparisons in modelled neurons and **(C)** recorded neurons (length scale σ = 2); error bars are standard deviation. **(D)** The PD map in a topo-VAE with σ = 2, the hyperparameter value which well approximates the recorded neuron topography (cf. Supplemental Fig. 3 for other values of length scale σ).

In the case of the model, different choices of the neighbourhood hyperparameter, σ, led to differences in the exponential distributions of PD difference (Supplemental Fig. S3.B,C). Small neighbourhood values (σ=1, Supplemental Fig. S3.B,C, left) led to more uniform distributions, whereas larger neighbourhood values (σ={2,3}, Supplemental Fig. 3.B,C, middle and right) led to exponential relationships like those we found in the recorded data, with the closest match to the model being σ = 2. Note, that for our simple cortical grid model we only considered integer values of σ.

Next, we examined which specific elements of the lateral interaction were important for reproducing the recorded data. We verified through model ablation the importance of the shape of the Mexican hat function (blue curve in Fig. 5.A), which determines the strength of excitatory and inhibitory connections as a function of distance between two neurons. We found no lateral effect function with a significant effect on input data reconstruction accuracy when compared to a model with no lateral effects (P<0.005; ANOVA). Flipping the function upside down (i.e., short range inhibition and intermediate range excitation, yellow curve in Fig. 5.A) eliminated the obvious topographic structure (Fig. 5.B, yellow box). Similarly, removing either the excitatory (purple curve in Fig. 5.A) or inhibitory component (red curve in Fig 5.A) produced results unlike the localised topographic structure of our primary model against which the recorded data was validated (Fig. 5.B, purple and red boxes, respectively), suggesting that the specific Mexican hat shape was important for reproducing biological results.

**Figure 5.**
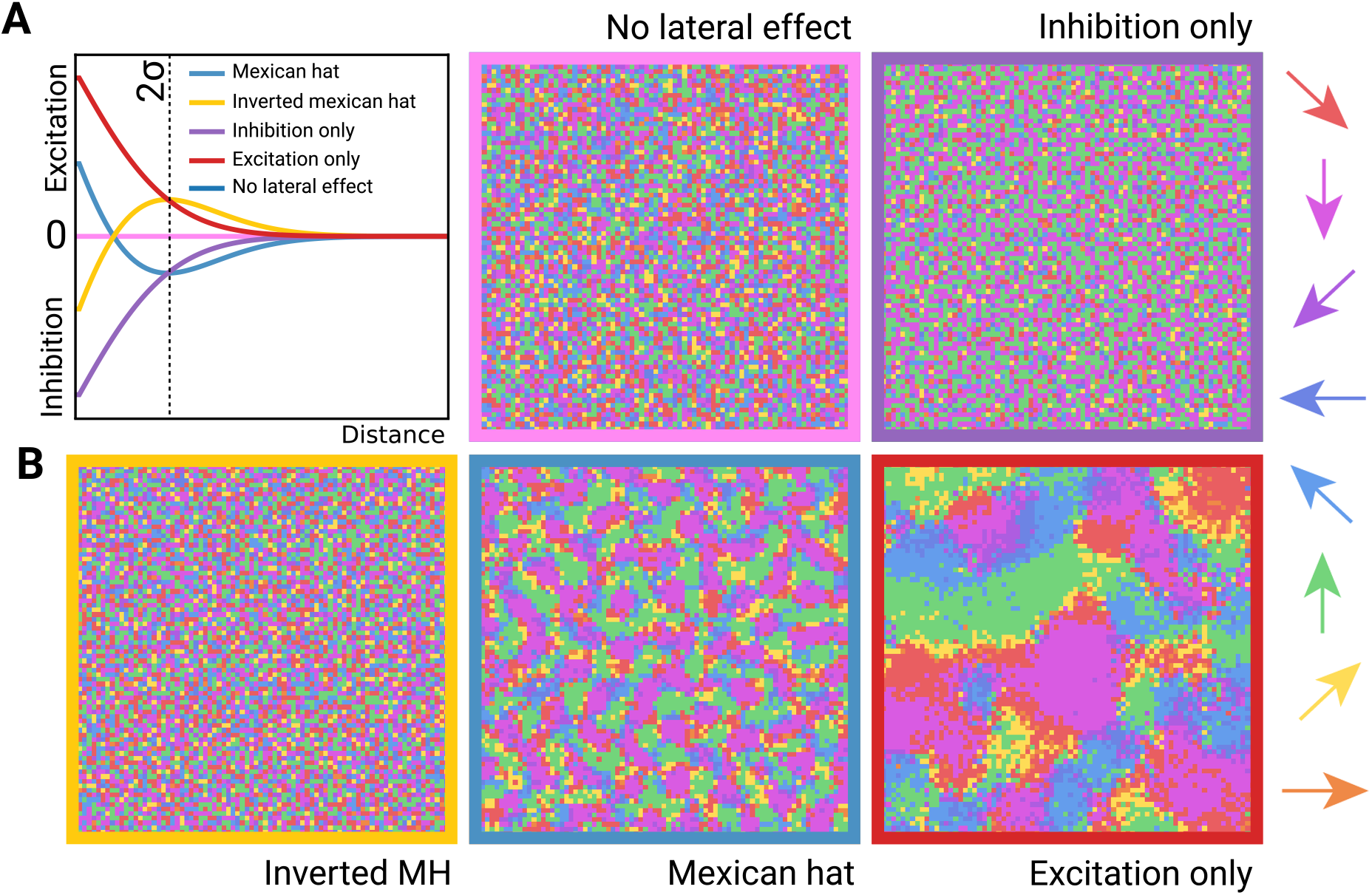
Mexican hat lateral effects are essential for reproducing spatial organisation. (**A**) Alternative lateral effect functions tested in control experiments, compared to the Mexican hat (blue line). **(B)** resulting PD maps for each function (colour matched to 5.D). The PD map for the inverted Mexican hat model is not shown due to space constraints, but produces no topographic structure, much like models with no lateral effect or inhibition only.

In addition to predicting the spatial organisation of proprioceptive coding, our model enables testable predictions about the representation of movements involving multiple joints (wrist, elbow, shoulder) across the neural population. To obtain a measure of how the 3 joints (Fig 6.A) are encoded across the cortical surface relative to each other, we found the average correlation with firing rate for all the degrees of freedom of each joint. All joints were evenly represented across the whole population (Fig 6.B), but no neurons correlated with only one joint. Many modelled neurons were correlated even with non-adjacent joints, e.g. elbow-wrist or even all three joints, possibly a result of the natural covariance patterns that exist between joints. For pairwise joint sensitivity comparisons see scatter plots (Fig. S3).

**Figure 6.**
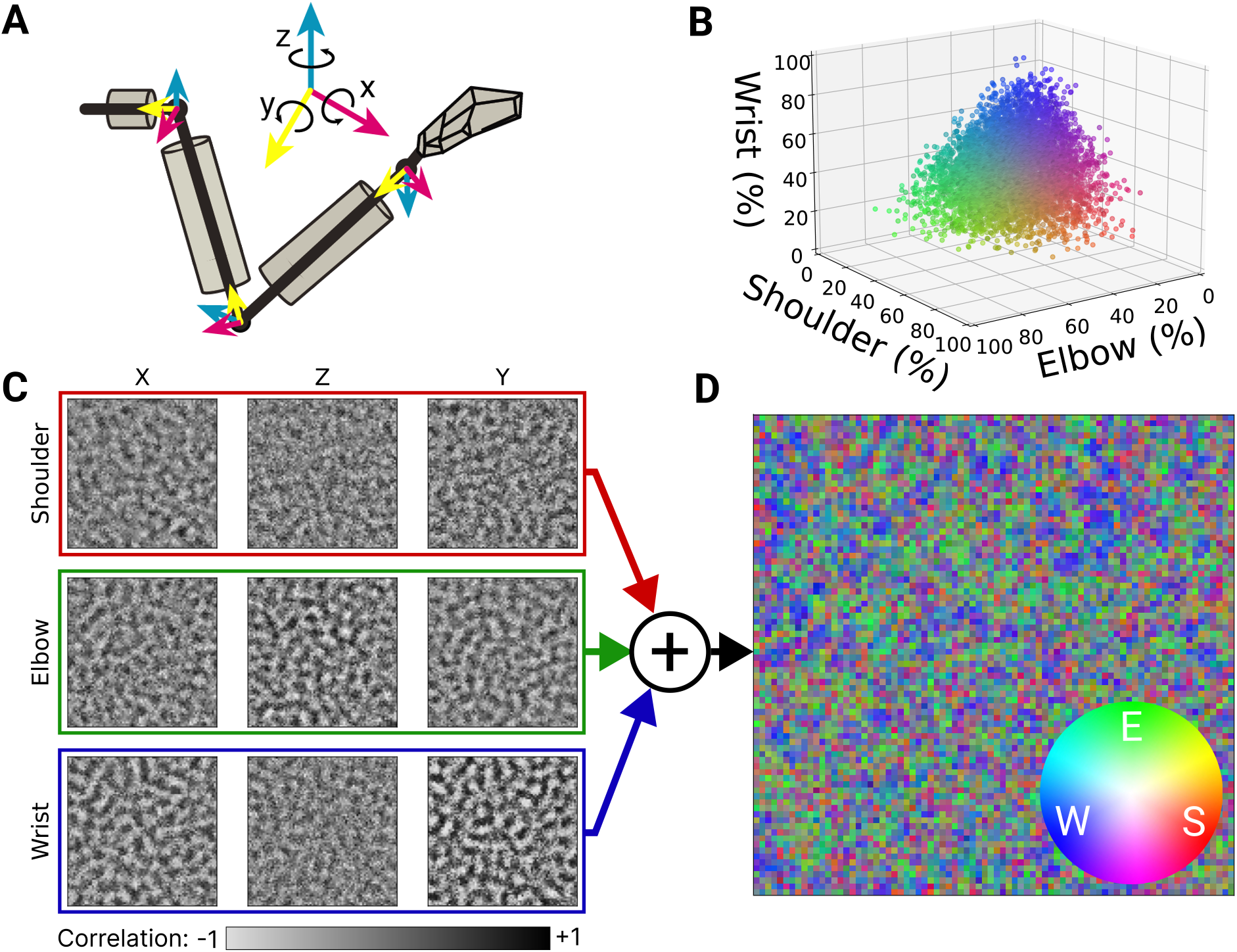
Sensitivity of latent layer neurons in topo-VAE to different input dimensions. (**A**) Joint angle axes, 3 per joint. **(B)** 3D plot showing the relative sensitivity of each neuron to joint inputs. Sensitivity is defined as the sum of correlations across X/Z/Y axes of all joints for a given neuron (colour normalised by maximum values per joint) (n=6400). **(C)** Correlations between each dimension of input joint motion and neural response. (**D)** Joint sensitivity map where each pixel represents a neuron and the pixels RGB values (colour wheel inset) are reflecting the correlation with shoulder (“S”), elbow (“E”), and wrist (“W”) angular velocity, respectively (colours normalised by maximum value across joints).

Fig. 6.C shows in more detail, the response strength of our cortical model neurons for individual degrees of freedom. The maps had a spatial pattern of ripple-like transitions between positive (white) and negative (black) correlations (Fig. 6.C), producing topographic clusters of neurons that are sensitive to particular joints (Fig. 6.D). The scale of these clusters approximately matched the that of the Mexican-hat lateral connection range. Regions of pure red, green, or blue in Fig. 6D would correspond to neurons responsive to only shoulder, elbow or wrist respectively. Instead, colours that are the result of blending two or more joints predominate.

## Discussion

Here, we combine natural proprioceptive stimuli with a novel model of neural coding and topographical organization of neurons in somatosensory cortex to help develop our understanding of proprioceptive coding and allowing us to paint a picture beyond that we can see through the restricted lens of neural recordings. Core to our approach was allowing the model to learn to represent the statistics of natural human movements rather than stereotyped movements, which under the efficient coding hypothesis is key to understanding sensory representations in the nervous system [23,29–31].

Our topo-VAE is designed to reflect information processing in cortex: we hypothesise that the cortex learns, through experience, an efficient representation of the somatosensory world which we replicate here as a deep autoencoder network. The network learns a nonlinear mapping from kinematic inputs into a “cortical” latent space of spiking neurons, which embodies a stochastic, generative model of movement-related neural activity. While VAEs have been used before to learn both single neuron and population level features from spike train data [32,33] a vanilla VAE (with a Euclidean latent space) cannot induce a topographic mapping on the data. Consequently, they cause nearby input variables (stimuli) to be represented in arbitrary locations across the latent space. By contrast, the latent space of a topo-VAE is arranged as a two-dimensional cortical surface, thereby creating a population of neurons on a square grid. This enables a lateral interaction term between neighbouring neurons to be added, that shapes cortical activity into local neighbourhoods, allowing it to learn cortical-like spatial structure as well as temporal single-neuron activity patterns, both of which we compare to recorded neural data.

Previous self-organising maps [19,34,35] operate on less biologically plausible grounds. For example, Kohonen-type maps operate on winner-takes-all, non-spiking activation (and consequently, their training updates synaptic connections in a winner-takes-all form as well) [35]. Therefore, only one neuron can ever be active for given a sensory stimulus in a Kohonen map, a property very unlike actual proprioceptive cortex. In contrast, any number of neurons in our topo-VAE can be active simultaneously (and consequently, all synaptic connections are updated as sensory information is processed). Likewise, Poisson GLMs consider only the input statistics when producing simulated neural responses. While it is possible to reproduce the coding properties of individual neurons from convergence of peripheral inputs to a GLM, those models can tell us little about the role of neighbouring cells in shaping receptive fields or neuroanatomy-neural function relationships in general.

Emergent from our model, were neurons with tuning surfaces comparable to those of recorded neurons as monkeys performed the same centre-out task (Fig. 2.A,B). Furthermore, higher-level features of the modelled neurons, such as PD distributions and the distributions of tuning curve features, were comparable to recorded neurons, particularly when the model was constrained by lateral effects and trained on stimuli that reflected the natural distributions of arm movement (Fig. 2&3). However, they were not captured perfectly (e.g. the PD and tuning geometry distributions of modelled neurons were still significantly different from those of recorded neurons, p<0.005; Kolmogorov-Smirnov test). Despite this, the current model is able provide multiple predictions about the nature of proprioceptive coding, one of which we partially validate with our limited neural data. Future work includes identifying additional constraints that can allow the model to better reproduce neural data from proprioceptive cortex.

The well-known somatosensory homunculus represents only the tactile component of somatosensation and its well-ordered map of the skin receptors. Since the human tactile homunculus has driven much of neuroscience’s intuition about somatosensory representations, it is tempting to hypothesise that proprioceptive representations might also be similarly structured. However, proprioception is driven not by receptors embedded in a “simple” two-dimensional sheet, but rather by different types of muscle receptors with varying dynamics, that span one, two, and even three joints. Thus, the expectation that proprioception and touch might share a similar homunculus may not be reasonable. Indeed, instead of a point-to-point mapping to the limb, in our model, we find structure akin to the pinwheels of the visual cortex, here representing hand movement direction instead of orientation selectivity. While the relatively large spacing of electrodes used to record neural data from area 2 does not allow us to confirm the pinwheel anatomical structure directly, there are signatures of it in the recordings, such as the fact that neurons recorded on a given electrode tend to have more similar PDs than those recorded on separate electrodes – a necessary, but not sufficient property for proprioceptive pinwheels.

Our model predicts not only the spatial organisation of tuning, but also the receptive fields of single neurons – although this prediction remains difficult to test with our currently available data. We find that modelled neurons encode combinations of joints, both adjacent (shoulder-elbow) and distant (shoulder-wrist). Such convergence in monkey somatosensory cortex has been observed for the hand [36,37] and must be present to some extent in the proximal arm, given its multi-articular muscles. Our model used only joint-based inputs (i.e., the kinematic state or pose of the arm) but knows nothing about the musculoskeletal mechanics of the limb (e.g., the fact that bi-articulate muscles span multiple joints). Nonetheless, multi-joint coding emerged for most neurons in our model. The correlations between motion of different joints that we observe in the natural data are substantial (4 principal components explain 80% of the variance of the seven degrees of freedom of the arm) and are the result of three main factors: 1) the biomechanics of the body, including the way many muscles span multiple joints, 2) the way the brain controls movements and 3) the tasks performed. Arguably, task requirements drive a substantial amount of the correlations. By training the topo-VAE on a highly varied dataset of natural movement, we enable it to generalise from these natural tasks to planar centre-out movements. Using this same dataset, Ejaz et al showed that the pairwise similarity of finger-specific fMRI activity patterns in human sensorimotor cortex was better explained by the correlation structure of hand movements than muscle activity [29]. Therefore, the emergence of multi-joint proprioceptive receptive fields may be a further example of the shaping of neural activity by higher-order features of movements [38], analogous to higher-order features of visual receptive fields, such as edges in V1 [18].

It is important to acknowledge that while the comparisons between modelled and recorded neurons are promising, it is largely a qualitative similarity, particularly at the level of population statistics. Understanding what further constraints might enable our model to better reproduce these features will be an important step in further elucidating the principles that shape proprioceptive representations in the brain. We also make the key assumption that the proprioceptive organizational principles captured by our topo-VAE model based on the kinematics of human movement would extend to monkeys. While this assumption seems to have largely held, there may still exist differences that contribute to the inconsistencies between our model and recorded neurons.

Another key limitation of the current model is the use of only joint kinematics. Biological inputs to proprioceptive cortex are received from muscle spindles and Golgi tendon organs, which convey muscle length, velocity, and force information. An important future direction will be to test the topo-VAE on such inputs, whether they be recorded experimentally or inferred from existing kinematic data using musculoskeletal simulations. Furthermore, because of its accessibility with Utah multielectrode arrays, we compared our modelled neurons to those recorded in area 2 of the somatosensory cortex, an area that combines cutaneous and muscle information in single neurons. Area 3a, on the other hand, has only muscle-receptor inputs, and shares many features with the adjacent motor cortex. One might expect an even closer correspondence between our modelled cortex and area 3.

Lastly, we would like to highlight the observation that, when trained on the stereotyped centre-out-data only, the topo-VAE poorly predicted recorded neuron data more poorly than when it was trained on the richer natural movement data. We interpret this to reflect the efficient coding hypothesis [22], whereby a population of sensory neurons are optimised to code stimuli representative of those found in their natural environment. Such codes are well documented in the visual [18] and auditory systems [39] – here we provide evidence that this may also be the case for proprioceptive neurons. This points towards the importance of working with training data that matches the natural distribution of stimuli when modelling sensory neurons, as it suggests important features of the neural code (such as the complex receptive fields predicted by our model) will only be elucidated under these conditions. This observation may be of great importance for the many proprioceptive and motor neurophysiology experiments that have been conducted in highly constrained lab settings, settings that may not contain adequate ethologically relevant kinematic statistics to uncover the true coding of cortical neurons. Undertaking electrophysiological experiments with a broader repertoire of movements may affect proprioceptive neuroscience as much as the adoption of natural images did for understanding vision.

## Methods

### Human Behaviour Scenarios and Natural Movement Data

We recorded full-body movements from 18 healthy right-handed participants in two experimental scenarios. In the natural behaviour scenario (see Fig. 1.B & Supplemental Fig. 1), subjects performed unconstrained daily tasks in a working kitchen environment. As food preparation and feeding are universal behaviours, the only direction given to subjects was to prepare and eat an omelette. For this modelling work we used only the arm movement data (including wrist but not digits). The average recording time across subjects was 22 minutes. In the second movement scenario, a subject performed planar centre-out reaches in a 20×20 cm horizontal task space (see Fig. 1.C) to mimic the movement data for the monkey task. The horizontal task space was aligned 20 cm below the subject’s shoulder and centred on the mid-line at 30 cm forward of the chest.

Arm movements in both scenarios were recorded at 60 Hz by an XSENS 3D motion tracking suit, a full-body sensor network based on inertial sensors. We used biomechanical models and fusion algorithms (including calibration and validation) to estimate joint angles. Fig. 1.A shows the biomechanical structure and coordinate system used in this paper. Arm movement datasets were formatted as time series of angles between segments following the International Society of Biomechanics (ISB) Euler angle extractions [25] in a ZXY coordinate. For the elbow and the wrist, angular rotations of Z, X and Y represent flexion/extension, abduction/adduction, and internal/external rotation, respectively, since the biomechanics of the human body cannot be fully described by a rotation around a single axis and must instead be described with respect to 3 rotational axes. During planar movements, we used optical tracking as well as the motion tracking suit for capturing the endpoint (hand) position on the task square. Data from the inertial and optical motion tracking systems were synchronised manually via cue-based movements before, during and after the recording period.

### Variational Autoencoder with Topographic Latent Space

In the following we lay out the rationale for building our model and the model itself, the various forms of data we collected for model training and validation, and the validation methodology.

The topo-VAE model (cf. Fig. 1.D-G) uses the autoencoder framework to model sensory representations. Autoencoders are a type of artificial neural network used to learn efficient encodings of unlabelled data [40]. The neuroscientific mechanism is referred to as the Infomax principle, i.e. unsupervised learning to discover structure in the sensory data by maximising the match between the inputs (in our case, the somatosensory world) and their neural representation [21,22]. We choose specifically a variational autoencoder [24] because we want the latent cortical layer to be able to capture the stochasticity and variability inherent to neural representations [41]. We modelled the latent neurons as Bernoulli processes using the Gumbel-max trick [42] where neuron firing is sampled from a categorial distribution of {0,1}. This forces the model to learn an encoding based on spike counts, which for an unchanging input will approximate the Poisson distributed statistics of biological neurons. This allowed us to use the same data analysis pipeline as for our recorded neural data. However, this simple VAE model would be devoid of any spatial relationships between the neurons. Therefore, we added a simple organisational mechanism that would link learning between neighbouring latent neurons, effectively implementing cortical lateral connectivity with short range excitation and longer-range inhibition. Thus, our enhanced VAE learns both receptive field tuning properties (in an unsupervised manner through deep feature learning) and establishes a topographic relationship between neurons.

Let 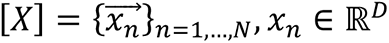 be the sensory stimuli, following the natural behaviour distribution 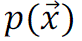. A group of cortical neurons, arranged with a topographic structure, are activated by the sensory stimuli [*X*] and generate firing patterns 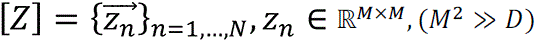. We aim to find a decoder 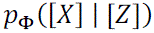 and corresponding neural responses [*Z*] to optimally represent the sensory stimuli [*X*], namely maximise the marginal likelihood of *p*([*X*]). The variational lower bound of the log likelihood log *p*(*X*) is derived as:

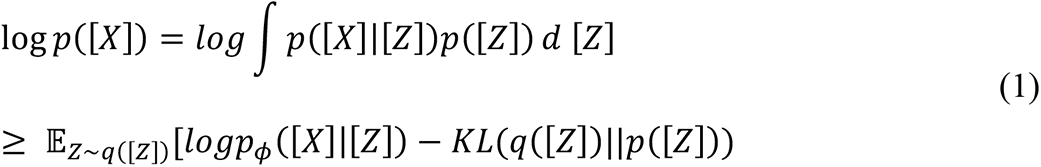

where *q*(*Z*) is the variational parameter approximating the intractable true posterior *p*(*Z*|*X*).

Our topo-VAE encoder can be considered as a multi-layer neuronal structure delivering sensory stimuli to the cortex from sensory afferents, through brainstem nuclei, to the thalamus and cortex. This implies that at the encoder level we do not attribute or consider specific representations at these intermediate stages of proprioceptive processing (including how different sensory systems are integrated. From the perspective of computational modelling, it is also the amortised inference for inferring the optimal representation *q*([*Z*]), helping to avoid smoothness problems in over-complete representations (‘back-constraint’). Fig. 1b illustrates the detailed structure of our topo-VAE, with a multi-layer perceptron (MLP) encoder and a linear decoder. The latent layer contains spiking neurons whose spike counts *Z*_n_ within a given time interval Δ*t* (i.e. over multiple input steps) follow a Poisson distribution. The encoder infers the distribution *q*([*Z*]) of responses for a group of cortical neurons, and *zn* is sampled from this distribution using the Gumbel-max trick:

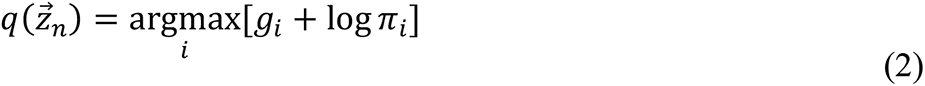

Where π_i_= [*p* 1 − *p*] is a vector defining the probability of a neuron spiking vs. not spiking, and *p* is set by the output of the encoder network for each neuron.

The decoder is a linear mapping from neural activities [*Z*] to the reconstructions 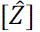 of the sensory stimuli. To summarise, the standard VAE model (which typically uses normally distributed random variables in the latent layer) is replaced by a latent layer of Bernoulli-type spike count distributions that are immediately applicable to neural signal analysis.

In addition to the log-likelihood function (Eq. 1) of the standard variational encoder, we include lateral effects in the latent layer. This was done by defining a distance-dependent function acting on the neurons, which are arranged in an *M* × *M* topographic map. A natural choice for cortical neurons is the Mexican-hat function, which transitions with distance from excitation to inhibition before vanishing [43] (Fig. 1.F,G). As we are modelling an entire population, we can represent the interaction between neurons as a matrix [Ψ]. Each element [Ψ*_p,q_*] represents the lateral effect between neurons p and q, calculated as:

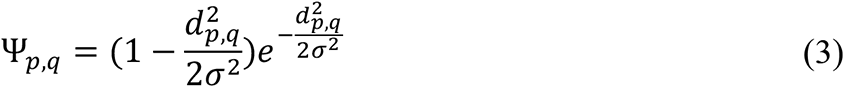

where *d_i,j_* represents the Euclidean distance and σ is a hyperparameter defining the common length scale of local excitation and intermediate-range inhibition. As shown in Fig. 1.G, the transition from maximum excitation to maximum inhibition spans a distance of 2σ and the lateral effect vanishes at about 4σ.

The total loss function governing our topo-VAE model is given by:

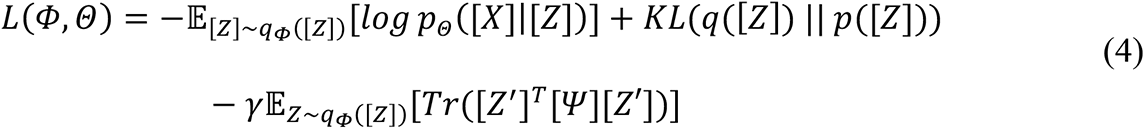

where *p*([*Z*]) is the prior distribution, set to be an independent Poisson distribution with rate r*_p_*, 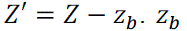 is the base firing rate and γ is a constant controlling the impact of the lateral effects. Φ and Θ represent the trainable parameters in the encoder and the decoder, respectively. The KL divergence term in the loss function performs as a constraint on temporal sparsity, penalizing firing rates far from the expected rate r*_p_* in the prior distribution *p*([*Z*]). r*_p_* is usually set to be small, due to constraints such as metabolic cost [44], and allows us to naturally control the temporal sparsity of neural activity. In addition to the temporal sparsity, our topo-VAE also involves structured spatial sparsity. The lateral effect, as represented by the topographic term 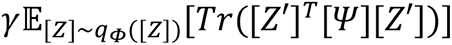 in Eq. 4 introduces topographic structure on the latent space and specifies the firing dependencies between neurons. Lateral excitation dominates the formation of pattern patterns in a structured space while lateral inhibition encourages spatial sparsity by penalising co-activation of non-nearby neurons. From the perspective of probabilistic inference, this regularisation item is equivalent to amending the prior distribution *p*([*Z*]) and modifies the target function as:

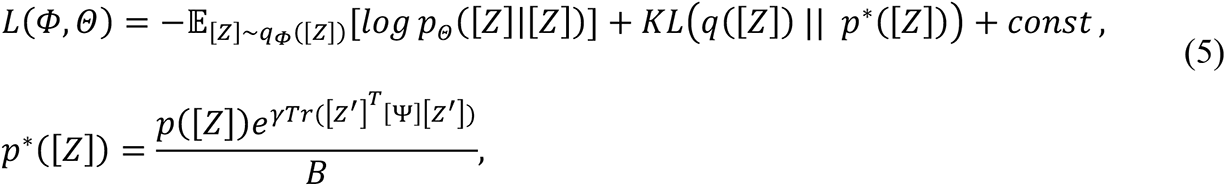

where the normalisation factor 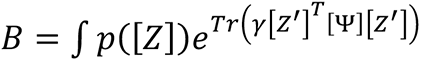 *dZ* is a constant. In this form, both the firing sparsity and the lateral effect are expressed within the amended prior distribution *q*^∗^([*Z*]) and the target function is maintained in a standard variational autoencoder framework.

We demonstrate through ablation and parameter variation experiments the need for the specific design elements of our model to explain the biological data.

The encoder component of the model contains two fully connected feed-forward layers of size 50 and 100 neurons, respectively, and a final linear readout layer. Tanh activation functions were used for all neurons in the first two layers. The reconstruction error of the model is measured using mean squared error.

To compare our modelling and recording results, we chose hyperparameters of the topographic map with consideration of 1) spatial densities of S1 neurons and 2) characteristics of the neural recording devices. The density of neurons in S1 is about 8M – 17M per cm^3^ [45,46] of which 70% to 90% are pyramidal cells [47]. The thickness of cortex varies from 1 mm to 4.5 mm [48,49]. The Utah electrode array has 100 microelectrodes arranged in a 10 x 10 configuration with 400 μm separation along each axis, thus spanning 3.6 mm x 3.6 mm of the cortex. We model neuronal anatomy as voxels or cubes, where the number of pyramidal cells contained within 1mm x 1mm x 1mm of cortex varies from 12,000 to 1,000,0000, which means every 1 mm along the cortical surface crosses about 20 – 50 pyramidal cells (see Fig. 1c). A surface-parallel slice captures a grid of 80 x 80 neurons. This conceptually simplified arrangement allows us to formulate a computationally tractable design of the latent layer neurons in topo-VAE. We can give a bit of intuition of our topographic model’s parameters to neuroanatomy: the range of the lateral effect parameter σ translates to about 1-2 neuron spacing on our cortical model grid (assuming a radius of dendritic input to these cortical neurons of around 200 μm [50]). Its precise value was determined pot-hoc using model selection by numerically sweeping for a range of σ and selecting the best fitting value.

To observe the effect of the neighbourhood range parameter σ on topography in the latent representation, we test values σ={1,2,3}. For the expected firing rate *p(x)*, we find stable topography around *p(x)*=0.01. The training loss function contains multiple components which can be given individual weightings. Here we use weightings of 10, 0.4, and 0.005 for the reconstruction, lateral effect, and firing sparsity components, respectively. In addition, we include an L2 norm on the model weights β=0.1. When sampling from the latent space during decoding, we parameterise the scaling of the rate parameter, to simulate sampling across different time windows. Varying this parameter in the range [0,1000] yielded optimal reconstruction at a scaling factor of 40.

### Model Training & Validation overview

The topo-VAE model was implemented in Python using PyTorch [51] and run on a GPU workstation. All models were trained for 4000 epochs using the Adam optimiser with a learning rate of 10^-5^ and a batch size of 400.

To train our topo-VAE we needed to pre-process the input data. Empirical distribution of joint angular velocities during movements in our tasks is symmetric, unimodal, with sharp peaks at zero and heavy tails towards large speeds. To ensure equal contributions to model optimisation, these features are standardised without mean centring (since they are already approximately zero-centred, and to preserve the correspondence of a zero-valued input to zero joint angle velocity). This pre-processing also improves the training efficiency and convergence of our model without breaking the spatial structure of natural movements. In addition, since the original kinematic data were sampled at a frequency of 60Hz, we subsampled the training data at a depth of 1%, to remove redundancy and improve training times, without significant effects on the outcome of the model.

Optimisation of VAEs by backpropagation requires estimation of parameter gradients with respect to loss. Since gradients cannot be estimated for the Gumbel-max function, we implemented the Gumbel-softmax reparameterisation trick [52] during training, where *z* is sampled from *q(z)* using the Gumbel-softmax function:

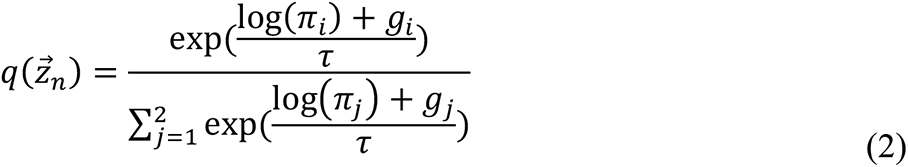

And *g*_i_ is a random sample from the Gumbel distribution *G*(0,1):

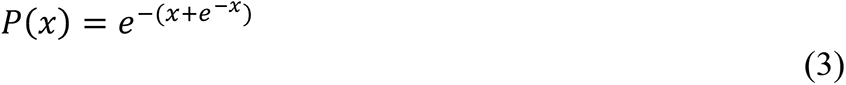

We wanted to perform feature learning of proprioceptive representations with our topo-VAE and this required large amounts of natural movement data. Therefore, we performed human experiments (see above) and measured the full-body kinematics of human subjects. We use human full-body behaviour as a functional proxy for non-human primate movements of the arm [53], as human data is much easier and more precisely obtainable. We used the human data to drive the training of our topo-VAE, then froze our model parameters to evaluate it. Our topo-VAE model is generative, so by playing back any limb movement data (time series of body poses) we obtain spike trains for each neuron in our cortical grid.

To compare our model’s predictions to those of recorded neurons, we use the kinematic data from humans performing the same centre-out task as the monkeys (see above) to drive the frozen topo-VAE model and compare its output to the actual recorded neural data.

### Model Robustness and Parameter Variation

The topographic property of the VAE arises from the lateral effect component of the loss function (Eq. 4). As a control, we tested our model with no lateral effects (Fig. 5.A-D), and with several different distance functions (Mexican hat, inverted Mexican hat, excitation only, inhibition only; Fig. 5.F,G). The inverted Mexican hat lateral effect is the additive inverse of the Mexican hat function (Eq. 3). Excitation-only lateral effect is defined as:

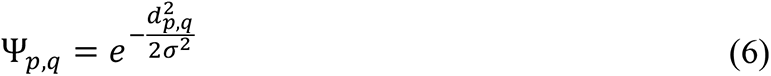

and the inhibition-only lateral effect is the additive inverse of the excitation-only lateral effect (Eq. 6).

### Model Analysis

To quantify the sensitivity of neurons to specific joints (Fig. 6), we individually perturb each input feature of the model and measure the Pearson correlation between individual neurons in the latent space and that feature. Since each joint is represented by three input features (angular velocity in the Z, X, and Y axes), we use the mean of the absolute correlation across all three axes to quantify the sensitivity of a given neuron to a particular joint.

We computed angular velocity profiles from the recorded data and analysed the natural movement dataset *X_nat_* and the planar movement dataset *X_pl_*. The first two principal components of the planar movements explained over 95% of the total variance, but only half of the variance of the natural movements. This reveals that joint velocities of planar movements are highly constrained, as expected, restricted largely to a 2-dimensional subspace. We applied the manipulative complexity metric [54] to quantify the complexity of the movements, which is defined as:

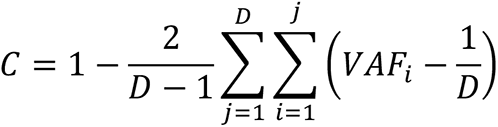

Where *VA*F*_i_* is the variance captured by the i^th^ PC. Larger values of C indicate higher complexity and a value of *C* = 1 means that all PCs contribute equally to the total variance. The complexity of planar movements was 0.06, much lower than the natural movement complexity of 0.5. We defined the direction θ and speed *v* of planar movements in world coordinates: 90°/270° are respectively away from and towards the chest, 180°/0° are to the left and right.

Fig. 1.H illustrates our use of the two human datasets with our computational model. The natural movement dataset *X_nat_* can be viewed as a group of samples generated from the natural movement statistic. Following the idea of natural sensory coding [18], we used this natural movement dataset as training data to learn an optimal neural coding scheme. After training, we tested the converged model with data from the planar reaching task. To compare with hand-based coding properties of area 2 neurons [55,56], we found a linear mapping between joint angular velocities and planar hand velocity. This allowed us to assess the relationship between hand movement direction and topo-VAE firing rates.

### Nonhuman Primate Behaviour and Data Collection

We used a combination of previously recorded data in which three rhesus macaques performed a planar, centre-out reaching task while seated, using a two-link planar manipulandum. A cursor displayed on a monitor tracked the position of the manipulandum and provided visual feedback for the monkey as he reached for a target on-screen. The monkey moved the cursor to a central target in the workspace. After a random delay period, 1 of 8 targets spaced evenly in a circle around the central target appeared on the screen and the monkey moved the cursor toward it upon an audible ‘go’ cue. After placing the cursor in the target for a random hold time of 0-500 ms, the monkey received a liquid reward and returned the cursor to the central target. We used 6 experimental sessions across three monkeys; two contain data that has been previously published [56], and the rest are unpublished. All procedures were in accordance with the Guide for the Care and Use of Laboratory Animals and were approved by the institutional animal care and use committee of Northwestern University under protocol #IS00000367.

Once a monkey was trained on the experimental apparatus, a 96-channel microelectrode array with 1 mm iridium-oxide coated electrodes (Blackrock Microsystems, Inc.) was pneumatically inserted in Brodmann’s area 2, near the intraparietal sulcus [56]. The implantation site was chosen to avoid cerebral vasculature and maximise proximal arm representation. All surgery was performed under isoflurane gas anaesthesia (1-2 percent) except during intraoperative recording to identify the arm representations, when the monkey was transitioned to a mixture of <0.5% isoflurane and remifentanil (0.4 ug/kg/min).

The data were recorded from the microelectrode array using the Cerebus multichannel data recording system (Blackrock Microsystems, Inc.). Thresholded waveforms and timing of behavioural task events were synchronised and recorded for offline analyses. The position of the handle was recorded at 1kHz. We discriminated single neurons using Offline Sorter (Plexon, Inc., Dallas TX). 383 recorded neurons were found to be suitable for subsequent analysis.

To calculate the preferred direction of a neuron, we used a simple bootstrapping procedure. For each iteration, we drew random points from the dataset conforming to a uniform distribution of movement directions. We then fit Poisson generalised linear models (GLM) with angular velocity inputs to the firing rate for the sampled timepoints. The GLM models are defined by:

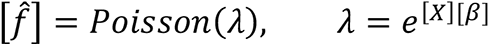

where 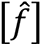 is a *T* (number of time points) by N (number of neurons) matrix of firing rate estimates of the recorded rates [*f*], *X* is a *T* by 2 (number of velocity inputs) matrix, and β is a P by N matrix of encoding parameters. β was found using maximum likelihood estimation. The preferred direction was then calculated from the encoding vector of the GLM as:

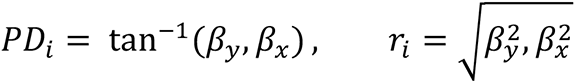

In these equations, a bootstrap PD estimate of neuron *i* was defined by β_y_ and β_x_, the bootstrap encoding parameters for hand velocity in the *y* and *x* directions, respectively. We took the circular mean of the PD estimates over all bootstrap iterations to find the PD for each neuron. This method of PD calculation was used for both recorded and modelled neurons.

## Data Availability

All code is available at https://doi.org/10.6084/m9.figshare.17393591.v3.

All data is available at https://doi.org/10.6084/m9.figshare.17393591.v3.

## Author contributions

AAF conceived the study. LEM & AAF supervised the study. YW, AAF developed the theory and model. JAH, AAF collected the behavioural data. KPB collected the unpublished neural data. YW, MG implemented the model code. YW, MG implemented the behavioural data analysis code. KPB implemented the neural data analysis code. KPB, MG, YW, AAF, LEM analysed the data. KPB, LEM, AAF drafted the manuscript. All authors discussed the results and reviewed the manuscript.

## Funders

We acknowledge: KPB was supported by NIH BRAIN NRSA Postdoctoral Fellowship F32MH120893 and [S1R01]. YW was supported by an NIHR Imperial College BRC Deep Phenotyping Grant and Project eNHANCE (http://www.enhance-motion.eu) under the European Union’s Horizon2020 research and innovation programme (Grant No. 644000). MG was supported by the Wellcome Trust PhD Program “Bioinformatics & Theoretical Systems Biology” (222888/Z/21/Z). JAH was supported by an EPSRC Doctoral Training Award. LEM was supported by NINDS grant #R01NS095251. AAF acknowledges his UKRI Turing AI Fellowship (EP/V025449/1).

The funders had no role in study design, data collection and analysis, decision to publish, or preparation of the manuscript.

## Financial Disclosures

MG, KB, YW, JAH, LEM, AAF have no competing financial interests.

## Notes

https://doi.org/10.6084/

### Competing Interest Statement

The authors have declared no competing interest.

### Summary of Updates

The work evolved further requiring to update the main text and the results. This include improvements to figure legends to improve clarity of number of model seeds tested, and model architecture. Further addition of supplemental figures outlining controls for spatial organisation constrains in latent space, hyper parameter tuning and sensitivity of model results to hyperparameters. Update to referencing style. Change of authorship order to account of ongoing effort by Max D Grogan on the manuscript as agreed by all authors.

