## Supplemental material for "Predicting proprioceptive cortical anatomy and neural coding with topographic autoencoders"

* Corresponding author

#### *Supplementary Figures*

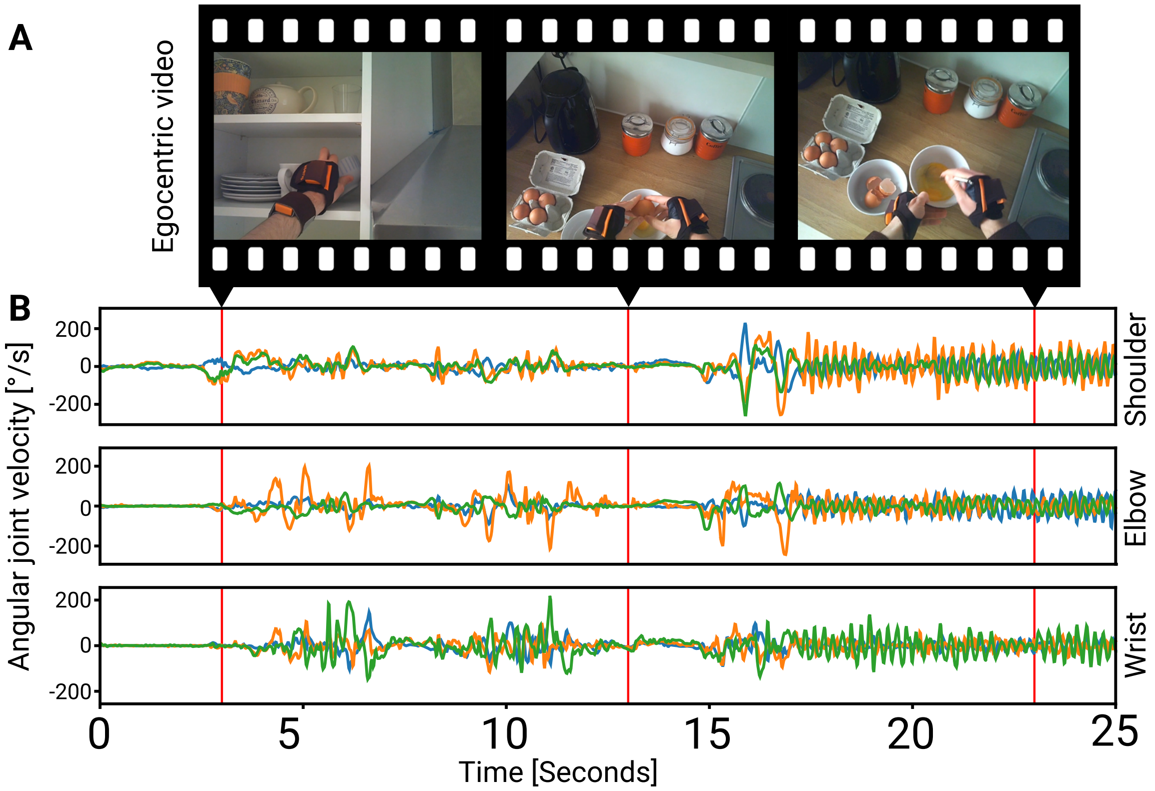


***Supplemental Figure 1.*** **Natural kinematics from behaviour in daily life.** ***(A)*** *Examples of natural behaviours contained in the dataset, as seen through egocentric video.* ***(B)*** *9-dimensional angular joint velocity time series of XZY joint axes (blue/green/orange lines, respectively) for shoulder, elbow, and wrist joint. Red lines correspond to the frames shown in (A).*

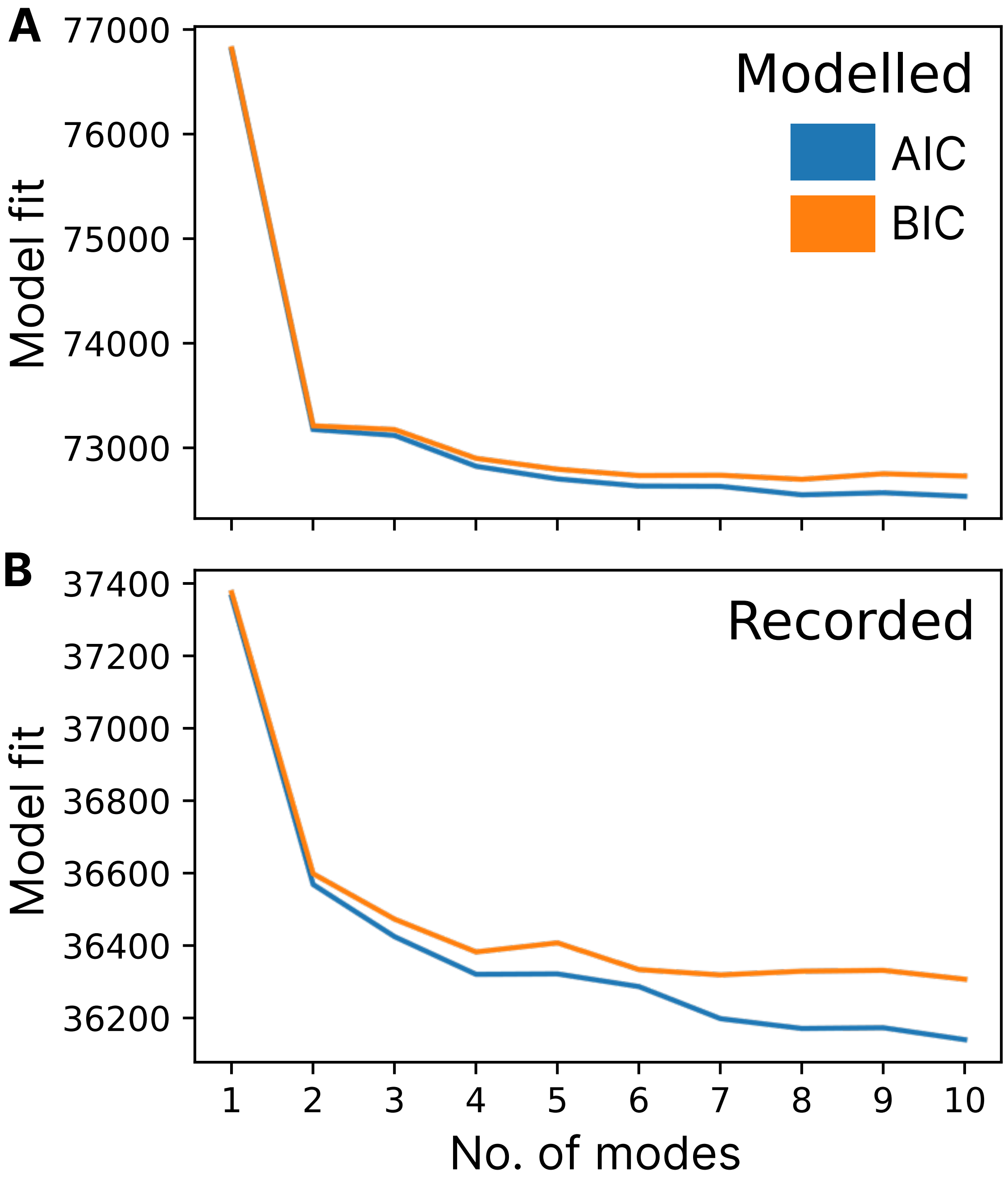


***Supplemental Figure 2.*** **Bimodality of PD distributions.** *Fit quality (AIC & BIC) of Gaussian mixture model fits with a varying number of modes for the preferred direction (PD) distributions of **(A)** Modelled neurons and **(B)** Recorded neurons. Note that model quality continues to improve (decreasing AIC/BIC value) with number of modes, however the most significant decrease occurs between 1 and 2 modes, while remaining improvements are small.*

*This suggests the PD distributions of both modelled and recorded neurons are well captured by a bimodal distribution.*

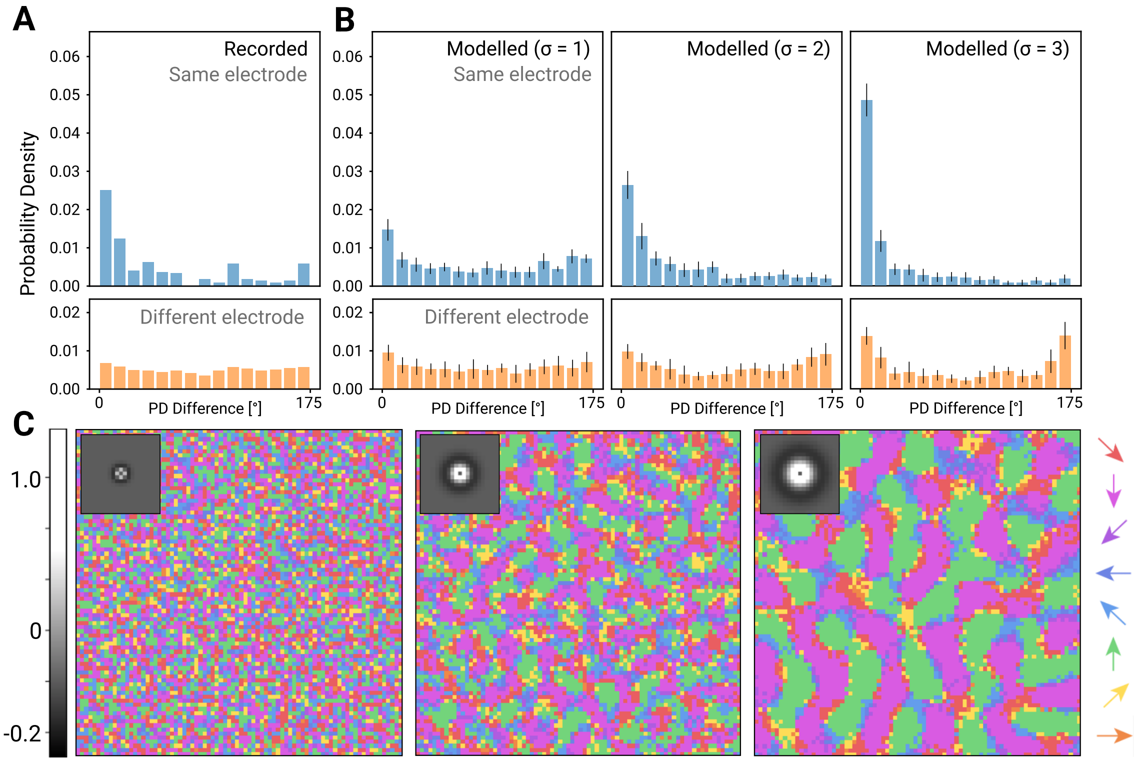


***Supplemental Figure 3.*** **Effect of σ on cortical topography.** ***(A)*** *Distribution of PD difference for same electrode and different electrode comparisons in recorded neurons and* ***(B)*** *modelled neurons, with σ=1, 2, and 3 shown on the left, middle, and right, respectively (n=6400). Standard deviation across 10 models with different weight initialisations is indicated by a black bar for each bin.* ***(C)*** *PD maps for σ=1, 2, and 3 shown on the left, middle, and right, respectively. A visualisation of the lateral effect under σ is also shown inset for each map.*

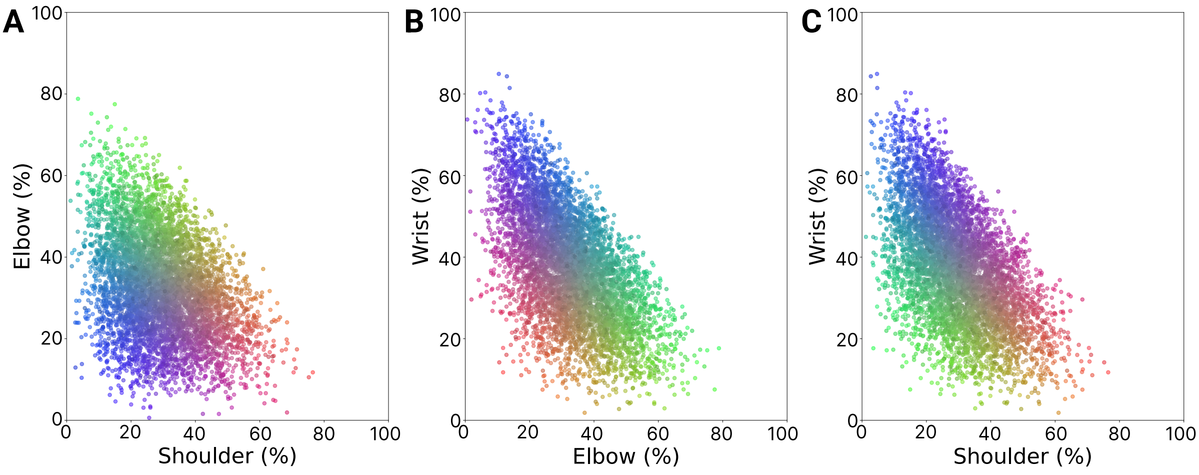


***Supplementary Figure 4.*** **Tuning of modelled neurons to arm joints.** *Tuning is quantified by the correlations between the activity of a neuron and joint angle velocity inputs from the shoulder, wrist, and elbow (summed across X/Z/Y planes for each joint). The sum of correlations across joints is then normalised such that a neuron with equal elbow, shoulder and wrist tuning would be positioned at 33.33% for each joint in the above plots. Each plot shows the relative tuning of each neuron (one data point) to joint inputs for the following pairs:* ***(A)*** *Elbow vs Shoulder,* ***(B)*** *Wrist vs Elbow, and* ***(C)*** *Wrist vs Shoulder. The colour of each dot is set by using their x, y, and z location in the plot to drive their respective red, green, and blue colour channels. The 3D plot of neurons for wrist, elbow and shoulder together are shown in Fig. 6 in the main text.*

### *Supplementary Methods*

An earlier version of this model was also developed in which the encoder network parameterises a Poisson distribution with rate parameter $\lambda$. In that model, the sampling step was not performed in a manner that allowed for gradient estimation with respect to reconstruction, meaning the model was optimised to decode inputs from a noisy, topographically organised, random projection of the original input. That model could still achieve moderate reconstruction performance, and both the spatial organisation and tuning of individual neurons was comparable to that observed in the full differentiable Bernoulli neuron model presented in the main paper (Fig. S4).

#### *Supplementary Code & Data*

The supplementary code and data for all main figures presented in this manuscript are found in a Figshare repository here: https://figshare.com/s/e028d90e33041bd5c3b3

The Figshare will be made publicly accessible upon publication.
